# Energy Crisis after Inter-System Mitochondria Transfer is the Direct Cause of Death by Sepsis

**DOI:** 10.1101/2022.09.01.505533

**Authors:** Michael Tang

## Abstract

Sepsis is one of the leading causes of death worldwide. With nearly 50 million incidences per year, it causes 11 million deaths worldwide annually, exceeding the 10 million total deaths caused by all tumors. Surprisingly, there is no specific drug available on market, explaining why it has a mortality rate as high as 22.5%. The lack of specific drug is mainly caused by the lack of understanding of how sepsis causes death. In this paper, I hypothesized that since energy production by mitochondria through respiration is not sustainable because the high level of reactive oxygen species (ROS) produced during respiration damages mitochondria themselves, mitochondria in the immune system cannot meet the dramatic and long-lasting high level of energy requirement of the system during sepsis. The immune system uses up all the functional mitochondria in the body by inter-system mitochondria transfer (ISMT), which dumps its used, unfunctional, or oxidized mitochondria to and recruits functional mitochondria from other systems. ISMT leads to the lack of functional mitochondria, hence energy, in the brain and the heart, and eventually causes death of the body. The hypothesis was supported by three key results: First, 2.5 hours after sepsis induction, mtDNA copy number increased dramatically in the spleen, brain, muscle, and blood, but decreased dramatically in the liver, kidneys, and skin. Second, mice died from sepsis showed a severe decline of mitochondria function in the brain and the heart. Finally, a single injection of isolated functional mitochondria to mice with sepsis reduced the mortality rate compared to mice received inactivated mitochondria.

## Introduction

Sepsis is the over-reactivation of the immune system to an infection. It is predominantly triggered by bacterial infection, but can also be triggered by virus, fungi, or parasite infection. The over-reactivation of the immune system damages the body, leading to multiple organ failure and may eventually causes death. Sepsis is one of the leading causes of death in the world. It was estimated that 48.9 million cases occurred and 11 million of sepsis-related deaths reported worldwide in 2017 (Rudd et al. 2020). This number is more than the 10 million total deaths caused by all tumors in 2021 (WHO 2022).

Surprisingly, the is no specific drug for sepsis. Treatment includes antibiotics, intravenous fluids, vasopressors and supportive care (Mayo 2021). Activated protein C was the first and only sepsis specific drug that has ever been approved. It was approved by FDA in 2000, but was withdraw around 2010 due to the lack of evidence that it is beneficial to patient (Lai and Thompson 2013). Due to the lack of specific treatment, the outcome of sepsis is dissatisfactory, with a mortality rate as high as 22.5%, 34-fold as high as that of COVID-19 in 2019 (Verity et al. 2020).

The lack of specific drug is caused by the poor understanding of the mechanism by which sepsis damages the body, causes organ failure, and eventually leads to death. Many mechanisms have been proposed to explain how sepsis damages the body, one of them is the mitochondria dysfunction. For example, it has been reported that sepsis mice or patient showed mitochondria dysfunction not only in the immune system (Li et al. 2010), but also in other organs or systems including the muscle (Welty-Wolf et al. 1996), liver (Aki et al. 2017), kidneys (Patil et al. 2014), heart (Dong et al. 1993, Lin et al. 2020, Stanzani et al. 2019), and the brain (Wu et al. 2015). However, sepsis is characterized by over-activation of the immune system, how over-activation of the immune system leads to mitochondria dysfunction in other organs or systems remains poorly understood.

During sepsis, energy requirement in the immune system increases dramatically and rapidly and usually lasts for days. Most of the energy is produced by respiration in mitochondria. Mitochondrion was derived from an ancient bacterium, and can transfer between cells both in vitro and in vivo. It has been known since 1982 that isolated mitochondria are able to enter surrounding cells (Clark and Shay 1982). After that, many studies reported that mitochondria can transfer between cells in vitro and in vivo (Ahmad et al. 2014, Chung-ha et al.2014, Hayakawa et al. 2016, Islam et al. 2012, Lou et al. 2012, Moschoi et al. 2016, Pasquier et al. 2013, Rebbeck et al. 2011, Rosina et al. 2022, Saha et al. 2022, Spees et al. 2006, Zheng et al. 2021). Based on the knowledge that mitochondria are able to transfer among cells, I hypothesized that energy crisis in vital organs such as the brain and the heart caused by inter-system mitochondria transfer (ISMT) is the direct cause of death by sepsis. The detailed hypothesis is as follows: since energy production by mitochondria through respiration is not sustainable because high level of ROS produced during respiration damages mitochondria themselves, mitochondria in the immune system cannot meet the system’s dramatic and long-lasting high level of energy requirement during sepsis. The immune system uses up all the functional mitochondria in the body by ISMT, which dumps its used, unfunctional, or oxidized mitochondria to and recruits functional mitochondria from other systems. ISMT leads to the lack of functional mitochondria, hence energy, in the brain and the heart, and eventually causes death of the body. To test this hypothesis, mtDNA copy number in different tissues, including the brain and the heart, in mice with or without sepsis was measured, and it was found that within 2.5 hours of sepsis induction, more than 2-fold of mtDNA copy number change happened, supporting the hypothesize that ISMT is triggered during sepsis. Mice died from sepsis had reduced mitochondria function in the brain or the heart compared to that of the control healthy mice. In addition, sepsis mice received a single injection of active, isolated mitochondria had reduced mortality rate compared to sepsis mice received inactivated, isolated mitochondria. All these results support the hypothesis that the death of sepsis is caused by lack of energy in vital organs following ISMT.

## Methods and materials

### Animals

All mice were obtained from TopBiotech and all procedures were reviewed and approved by the Animal Care and Ethics Committee of TopBiotech.

### mtDNA copy number

5 pairs of male Kunming mice (20-26 g) were divided into two groups randomly. The body weight difference was no greater than 1 g for each pair. Each mouse in sepsis group received intraperitoneally injection of 30mg/ml DH5-alpha E. coli in phosphate-buffered saline (PBS) at 0.1 ml/10g. Each mouse in the control group received intraperitoneally injection of PBS at 0.1ml/10g. 2.5 hours after sepsis induction while symptoms occurred, mice were sacrificed by cervical dislocation. After abdominal hair was removed by a pet shaver, 75% ethanol was used to disinfect the whole body for 3 min. Then excess ethanol was removed by dry paper towel.

After that, the abdominal cavity and chest cavity were opened. 40 μL of blood was collected from the heart in the presence of 1 μL of low molecular weight heparin and 20 mg of tissue from the heart, liver, spleen, kidney, brain, abdominal skin, muscle of the right shaft, fat of the mesentery was collected from each mouse. The blood or the same tissue from all the five mice in the control or sepsis group were combined and stored at −80c immediately before DNA isolation. DNA isolated by QIAamp DNA Mini Kit (Qiagen) was used as the template for qPCR. mB2MF (F: ATGGGAAGCCGAACATACTG, R: CAGTCTCAGTGGGGGTGAAT) and mMito (F:CTAGAAACCCCGAAACCAAA, R: CCAGCTATCACCAAGCTCGT) were used as the primers for nuclear DNA and mtDNA respectively. The 20 μL PCR reaction includes 2 μL of DNA template (20 ng/ μL), 0.4 μL of each primer (10 μM), 2x SYBR Green PCR Master Mix (AceQTMqPCR SYBR^®^ Green Master Mix, Vazyme, China), and 7.2 μL of dH2O. qPCR was performed on a Applied Biosystems ViiA 7TM Real-Time PCR System (ABI, USA). The reaction was initiated at 95°C for 5min, followed by 45 cycles through 95°C x 15s, 62°C x 60s, then a final cycle of 95°C x 15s, 60°C x 60s, and 95°C x 15s. All reactions were run in triplicate. Amplification curves were analyzed using Applied Biosystems ViiA 7 Software. The results were exported as Excel files, and Excel 2019 was used to calculated the mtDNA/nDNA ratio in each sample.

### Mitochondria isolation and inactivation

Mitochondria were isolated from either neonatal mice or adult mice. For mitochondria isolated from neonatal mice, postnatal day 1-3 mouse pups were disinfected with 75% ethanol before euthanasia by decapitation. The body was then cut into pieces and transferred to an ice-cold 200 ml glass bottle to be prepared for homogenization. For mitochondria isolated from adult mice, male or female Kunming mice (20-36g) were disinfected by 75% ethanol before sacrificed by cervical dislocation. The abdominal cavity and chest cavity were then opened. The heart, liver, spleen, and kidneys were removed and transferred to an ice-cold 200 ml glass bottle. 4 ml of ice-cold PBS, 0.04 ml of 0.5 M EDTA (PH 8.0) and 4μl of 200 mM phenylmethylsulfonyl fluoride (PMSF, dissolved in isopropanol) were added to each gram of tissue. The tissues were then homogenized using an electric homogenizer at the speed of 6000 rpm for 2 min. After that, the homogenized tissues were transferred to 50 ml tubes and were centrifuged at 1000 g and 4 °C for 10 min. The supernatant was then transferred to new 50 ml tubes, centrifuged at 12000 g and 4 °C for 10min. Afterwards, the supernatant was removed and the pellet was the isolated mitochondria.

For inactivation, isolated mitochondria were resuspended in PBS at the final concentration of 25% (weight to volume). Then, 0.5 ml of 75% ethanol was added to every 1 ml of mitochondria suspension and mixed well. After incubated at room temperature for 5 min, they were centrifuged at 12000g for 10min. The supernatant was removed, another 30 ml of PBS was added to each tube, resuspended well, and let sit at room temperature for 10min. After that, they were centrifuged again at 12000g for 10 min. Afterwards, the supernatant was removed and the pellet was the inactivated mitochondria.

### Mitochondria respiration

4 pairs of 6–8-week-old male Kunming mice were divided into two groups randomly. For each pair, the body weight difference was no greater than 1 g. Each mouse in the sepsis group received an intraperitoneally injection of 60mg/ml DH5-alpha E. coli in PBS at 0.1 ml/10g, while the control group received an intraperitoneally injection of PBS at 0.1ml/10g. All mice in the sepsis group died within 6 hours after sepsis induction. When one mouse in the sepsis group died, the corresponding mouse in the control group was sacrificed by cervical dislocation. Then, mitochondria of the heart or the brain of each mouse were isolated from each mouse by the way mentioned above.

After that, 50mg of isolated mitochondria were resuspended in 10 ml of mitochondria respiration buffer (10 mM HEPES, pH 7.4, containing 250 mM sucrose, 1 mM ATP, 2.24 mM ADP, 9 mM sodium pyruvate, 5 mM sodium succinate, 2 mM K2HPO4, and 1 mM DTT) in a 50 ml centrifugation tube. A drop of vegetable oil was added to the surface of the mitochondria suspension in each tube to insulate oxygen from the buffer. The oxygen level was measured using a dissolved oxygen meter immediately (dO2 0min) and 30 min later (dO2 30min). Mitochondria respiration rate was calculated by (dO2 30min-dO2 0min)/30 min.

### Sepsis induction and treatment

12 pairs of male Kunming mice (20-36 g) were divided into two groups randomly. The body weight difference was no greater than 1 g for each pair. 20% E. coli stored at −80c were thawed and diluted with PBS to the final concentration of 20 mg/ml. Each mouse received an intraperitoneally injection of diluted E. coli in PBS at 0.1 ml/10g. 2 hrs. after injection when the symptoms appeared, isolated mitochondria diluted in cell culture media Dulbecco’s Modified Eagle Medium (DMEM) supplemented with Penicillin/Streptomycin/Amphotericin B (final concentration was 700U/ml, 700μg/ml, and 1.75μg/ml respectively) were injected intravenously (IV) at the concentration of 25mg/ml and at the dose of 0.05ml/10g. The treatment mice received IV of active mitochondria and the control mice received IV of inactivated mitochondria. The death was recorded 1, 2, 3, 4, 5, 6, 9, 12, 24 hrs. and 2, 3, 4, 5, 6, 7 days after sepsis induction.

## Results

### Fast inter-system mitochondria transfer during sepsis

mtDNA copy number increased slightly in the heart and fat, increased dramatically in the spleen, brain, muscle and blood of sepsis mice compared to that of the control mice (Figure 1.). The change in the brain was so dramatic that it almost increased 10-fold. It decreased slightly in the skin, and dramatically in the liver and kidneys (Figure 1.). It is unlikely that the change is due to fast mtDNA replication and degradation since the change happened within only 2-3 hours. Meanwhile, while there are increase in some tissues, there are decrease in other tissues. Taken together, these results strongly indicate that there was transfer of mtDNA from the liver, kidney, skin to the heart, spleen, brain, fat, muscle, and blood, or from the digestive system and urinary system to the nerves system, circulatory system, immune system, and motor system.

**Figure 1.**
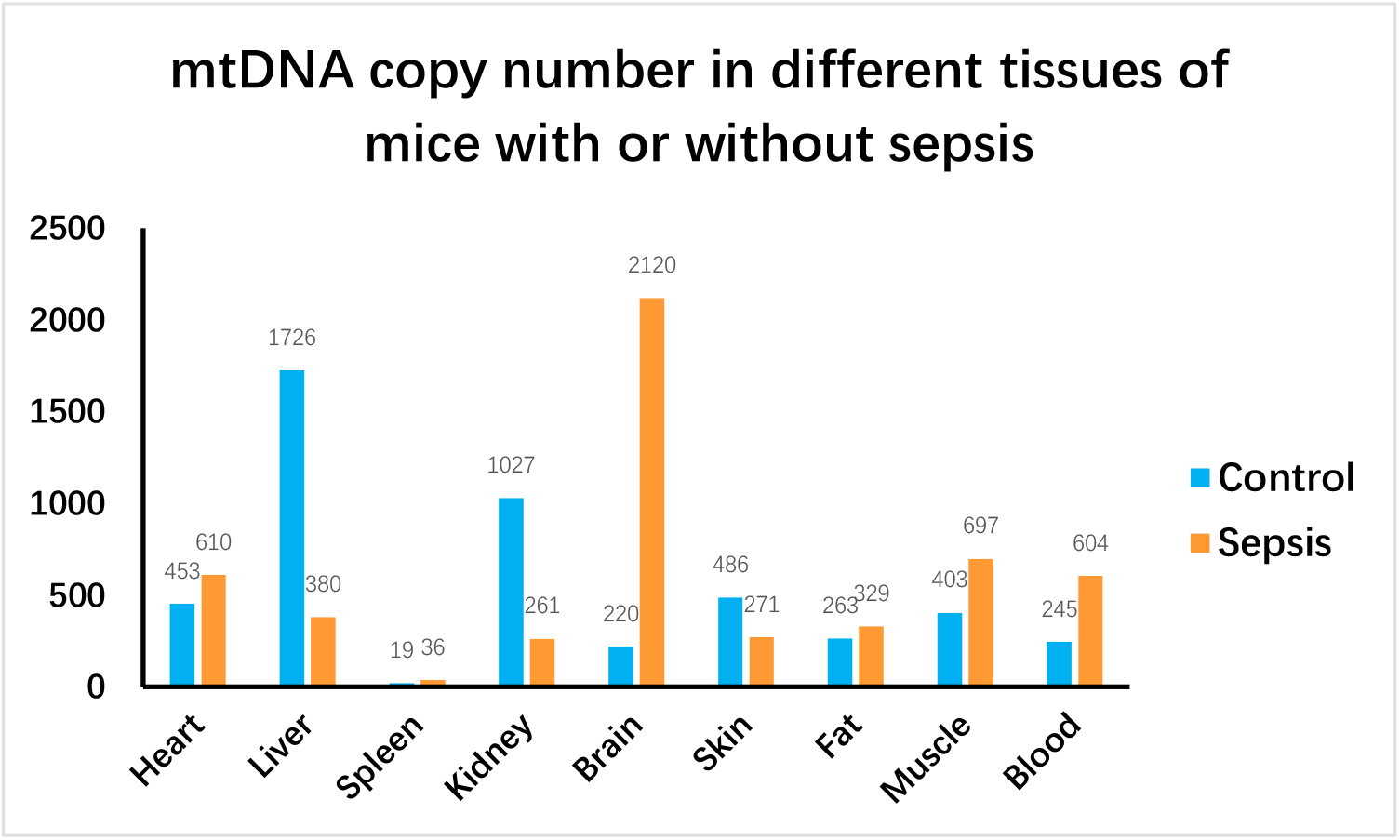
mtDNA copy number in the heart, liver, spleen, kidney, brain, skin, fat, muscle, and blood in mice with or without sepsis.

### Mitochondria function declined in the brain and heart

We then tested the function of mitochondria in the brain and the heart. Mitochondria were isolated from the heart, liver, spleen, kidneys, or brain of mouse died from sepsis or the control mouse, then oxygen consumption rate was tested. Consistent with our expectation, there was approximately 50% decline of oxygen consumption rate of mitochondria isolated from the heart or brain of mouse died from sepsis compared to that of the control mouse (Figure 2.). Meanwhile, oxygen consumption rate in other organs, including the liver, spleen, and kidneys didn’t change statistically significant (Figure 2.). In view of the high requirement of energy in the brain and heart, 50% reduction of energy production can be lethal.

**Figure 2.**
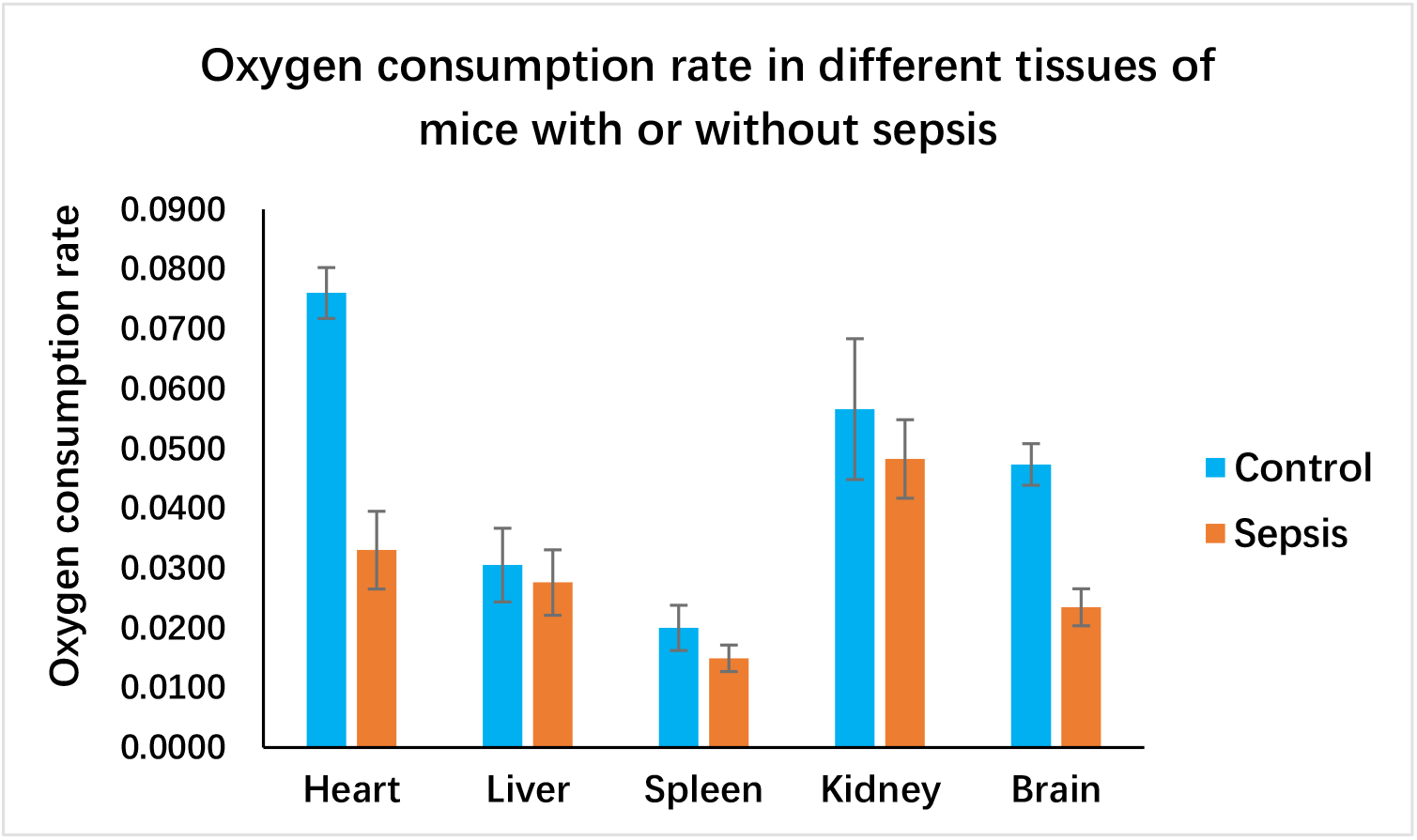
Oxygen consumption rate of mitochondria from the heart, liver, spleen, kidney, or brain of mice with or without sepsis. Error bars show standard deviation.

### Mitochondria transplantation increased survival rate of sepsis in mice

If energy depletion in the brain and the heart is the direct cause of death in sepsis, administration of active mitochondria should be able to reduce the mortality rate. To test this, we isolated mitochondria from neonatal mice, because their mitochondria should be of higher quality in theory. A single IV of 2.5% active mitochondria reduced the mortality rate for about 17% compare to mice received inactivated mitochondria after 1 week of observation (Figure 3.).

**Figure 3.**
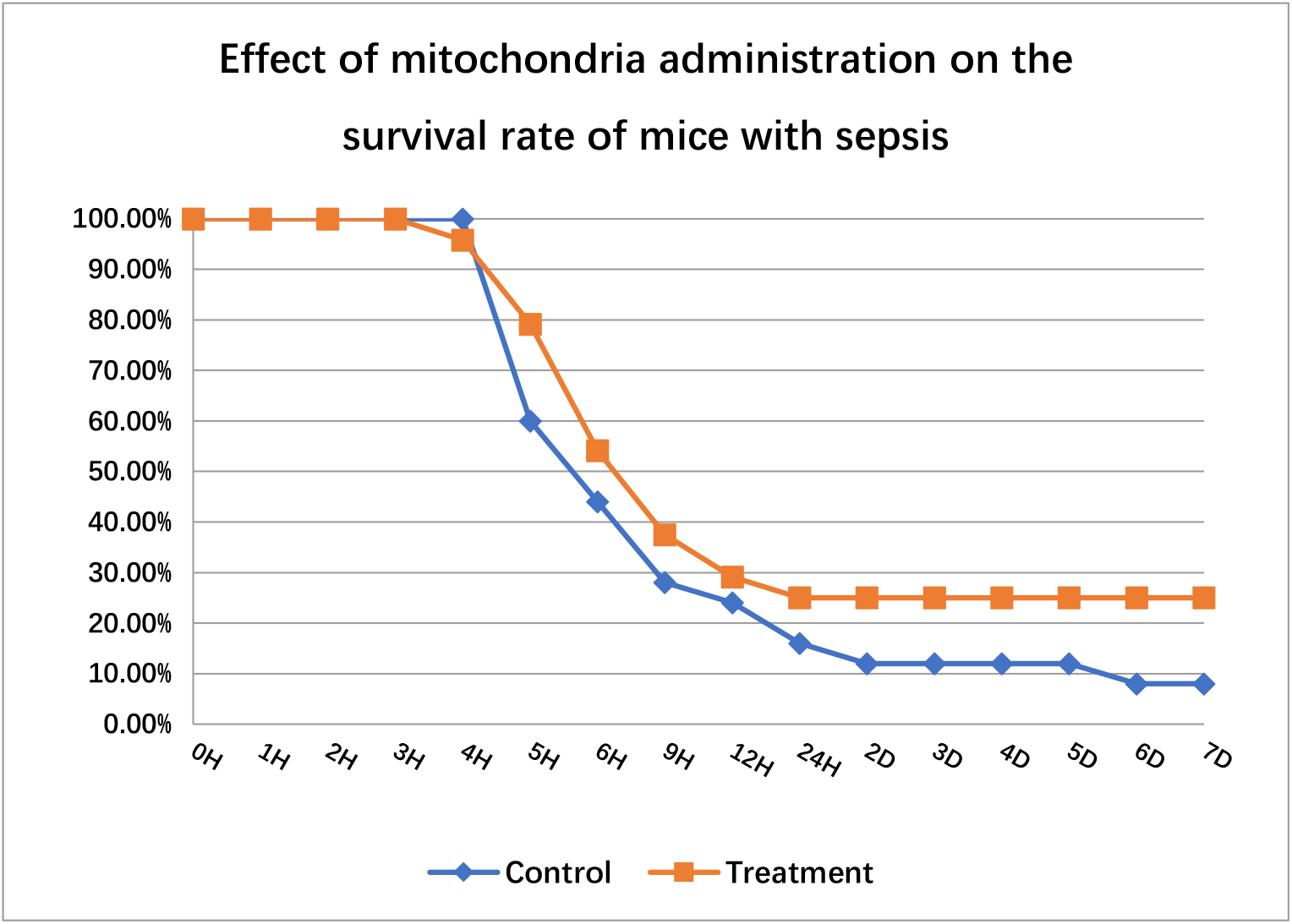
Functional mitochondria transplantation reduced the mortality rate of mice with sepsis.

## Discussion

The hypothesis and results will be discussed in the future.

